# Touch inhibits cold: non-contact cooling reveals a novel thermotactile gating mechanism

**DOI:** 10.1101/2024.08.06.606653

**Authors:** Ivan Ezquerra-Romano, Maansib Chowdhury, Patrick Haggard

## Abstract

Skin stimuli reach the brain via multiple neural channels specific for different stimulus types. These channels interact in the spinal cord, typically through inhibition. Interchannel interactions can be investigated by selectively stimulating one channel and comparing the sensations that result when another sensory channel is or is not concurrently stimulated. Applying this logic to thermal-mechanical interactions proves difficult, because most existing thermal stimulators involve skin contact. We used a novel non-tactile stimulator for focal cooling (9mm^2^) by using thermal imaging of skin temperature as a feedback signal to regulate exposure to a dry ice source. We could then investigate how touch modulates cold sensation by delivering cooling to the human hand dorsum in either the presence or absence of light touch. Across three signal detection experiments, we found that sensitivity to cooling was significantly reduced by touch. This reduction was specific to touch, since it did not occur when presenting auditory signals instead of the tactile input, making explanations based on distraction or attention unlikely. Our findings suggest that touch inhibits cold perception, recalling interactions of touch and pain previously described by Pain Gate Theory. We show, for the first time, a thermotactile gating mechanism between mechanical and cooling signals.

## 1. Introduction

The neural pathways that conduct information about a specific stimulus type from the skin to the brain are considered distinct somatosensory channels. These channels are thought to interact, for example by inhibitory synaptic connections in the spinal cord [1-5], and also supraspinally [6, 7]. For instance, touch reduces pain, and pain relieves itch [1-5, 8-10]. To study these interactions, researchers have selectively stimulated a target sensory channel and compare either neural responses or reported sensations when another sensory channel either is or is not stimulated. This research strategy has remained elusive for cold sensation because most cold stimulation devices inevitably require contact with the skin. Possible interactions between cold and touch could therefore only be investigated with controllable non-tactile stimulators [11].

Pain gating studies have shown that touch inhibits pain [1-3]. Different subpopulations of Aδ-fibres are thought to mediate both non-noxious cooling, and also heat pain in humans [2, 3, 12-17]. Additionally, recent studies have found robust and overlapping responses to both mechanical and cooling inputs in the mouse primary somatosensory cortex (SI) [18, 19]. In humans, SI BOLD activity can discriminate between warm and cold thermotactile stimuli applied to the hand [20]. Altogether, these results suggest that non-noxious cold may interact with tactile signals, for example, through gating mechanisms analogous to those previously reported for nociceptor signals.

Green and colleagues have reported that touch attenuates cold sensations in humans [21, 22]. They found more intense cold sensations when *making* tactile contact with an object already pre-cooled, a scenario they called dynamic touch, compared to when *maintaining* tactile contact with a thermally-neutral object that is then cooled to the same temperature, which they called static touch. However, both conditions in this study involved some degree of tactile input. In other words, skin cooling was not fully dissociated from touch. Understanding how touch modulates cold sensation would ideally involve comparing cold sensations with and without touch.

We have therefore studied detection of focal cooling with and without tactile stimulation, by using a novel non-tactile cooling stimulator [11]. We found that touch consistently decreased sensitivity to non-tactile cooling, recalling the interaction of touch and pain described by Pain Gate Theory [1-3].

## 2. Material and methods

### (a) Participants and ethics

A total of 36 healthy volunteers participated with ethical permission, 12 in each of 3 experiments (Experiment 1: 9 females, mean age: 25.92 years ± 5.57 SD; Experiment 2: 9 females, mean age: 28.33 years ± 6.74 SD; 9 females; Experiment 3: 11 females, mean age: 25.5 years ± 5.88 SD). The sample size was determined by a power calculation, as follows. We estimated an effect size of 0.857 (Cohen’s d) for the effect of touch on sensitivity to cooling, based on a previous study using a similar experimental design but showing that touch reduced sensitivity to pain [2]. For a one-tailed t-test, a significance level of 0.050, and a power level of 0.80, ten participants are required, but we decided to test 12 participants, for comparability with previous studies [2]. We defined *a priori* criteria to avoid floor and ceiling effects: overall response accuracy above 95% or below 50% in any condition would entail excluding the participant. In fact, no participant was excluded.

The research was approved by UCL Research Ethics Committee (ID number: ICN-PH-PWB-0847/010), and specific risk management protocols were approved and implemented with respect to thermal stimulation.

### (b) Experimental set-up

The experimental apparatus was similar to that described in [omitted for blind review] (figures 1a & b).

**Figure 1.**
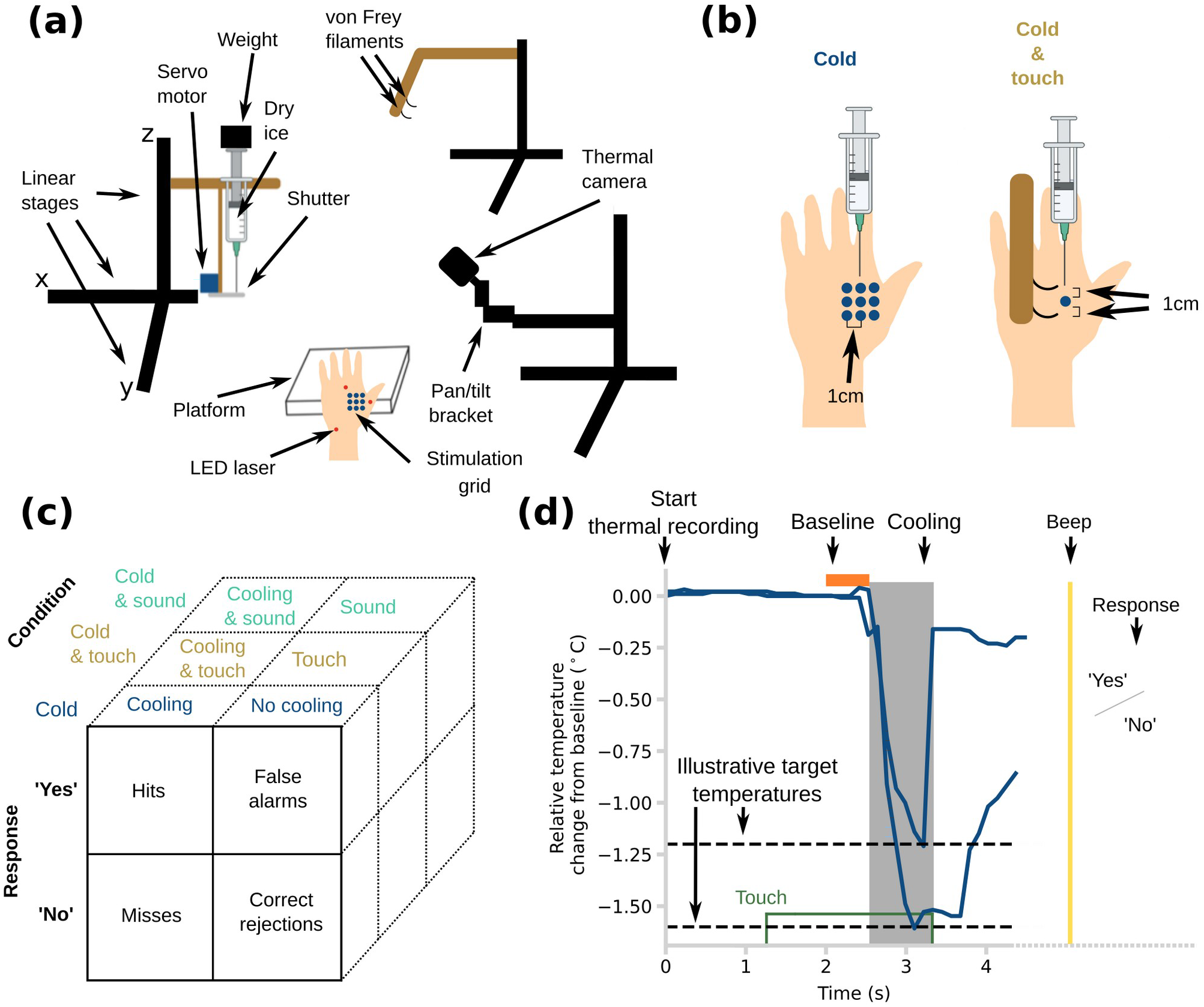
Experimental set-up, trial structure and design. **(a)** An illustration of the set-up with the main components including the mechanical stimulator. **(b)** Comparison between ‘Cold’ and ‘Cold & touch’ conditions **(c)** Table showing definitions of hits, misses, false alarms, and correct rejections for each trial type and condition based on response data. **(d)** Schematic displaying events on each trial including 2 illustrative thermal traces, with different target temperatures and from different participants. The traces show change of temperature from the mean of a baseline period of 0.5 s immediately before the thermal onset/tone onset. The grey shaded area indicates the period of thermal exposure which was accompanied throughout by a tone.

A tactile stimulus was delivered by two von Frey monofilaments [23] (bending force: 1 gram-force, diameter: 0.4 mm, length: 15 mm), aligned proximodistally, and each 1 cm from the cooling point (figures 1a & b). The position of the monofilaments was controlled by a computerised XYZ stage (Zaber Technologies Inc.) (figure 1).

Thermal tactile stimuli were delivered to the back of the left hand using a custom stimulator allowing controlled exposure to a small dry ice source. Nine skin locations, forming a 3×3 grid with 1 cm spacing, were thermally stimulated in pseudorandom order. The same location was restimulated with cooling only after at least 3 other locations had been visited, ensuring a minimum of 30 s for thermal recovery at each site between cooling events. On each trial, the distance between the dry ice nozzle and the skin was chosen based on the desired skin temperature decrease, using calibration values from a previous study [11]. A thermal camera on a pan/tilt head and additional XYZ stage (ROB-14391, SparkFun Electronics) viewed the stimulated skin region, and measured the actual temperature decrease on each trial.

To standardise skin temperature across participants and minimise variation in baseline skin temperatures, an infrared lamp (Infrasec IR2 250W bulb, Tungsram) controlled by a dimmer was used to gently warm participants’ hands at the beginning and during breaks. Windows and doors were closed to minimise airflows and thermal fluctuations in the room. A curtain blocked the participant’s view of hand and all apparatus.

The experiments followed a signal detection theory paradigm. In each trial, participants judged whether a temperature change was or was not present. In Experiments 1 and 2, a speech recognition algorithm was used to transform the participants’ responses (either ‘Yes’ or ‘No’) from voice to text (IBM Watson, IBM). Vocal responding was chosen because pandemic management protocols in place at the time mandated minimising manual contacts with surfaces. In Experiment 3, participants pressed ‘Yes’ and ‘No’ keys on a keypad (Pauk10, Targus International LLC).

In Experiments 1 and 2, the duration of possible stimulation was indicated with a tone (frequency: 400 Hz). In Experiment 3, the duration of possible stimulation was indicated by an LED (red LEDs, VCC) placed on the curtain between the participant’s eyes and the stimulated hand. The LED light was controlled by an Arduino Uno. In this experiment, some conditions involved auditory stimuli accompanying thermal stimuli. These were tones with a frequency of 500 Hz, a loudness of 50 dB at the position of the participant, delivered from micro-loudspeakers bracketing the thermal location. The aim of this experiment was to show whether the reduction in sensitivity to cooling was specific to touch or might also involve general factors such as distraction by any ongoing stimulus. We set the auditory intensity to be five times reported auditory threshold values [i.e., 10dB at 500 Hz; 24], since our tactile stimuli were also approximately five times previously reported detection threshold values of 0.2 gf [25].

### (c) Experimental design and task

At the beginning of all experiments, there were 4 training trials to familiarise participants with the trial structure and the task (2 cooling and 2 no cooling trials). Participant were instructed to focus on the thermal stimulus and respond ‘Yes’ or ‘No’ after a beep to the question: ‘Was there a temperature change during the tone?’. The question was presented after each stimulation by either a computer-generated voice (Experiments 1 & 2) or on-screen text (Experiment 3).

Each experiment involved an initial staircase to select stimulation levels, followed by a signal detection paradigm. A broadly similar exposure protocol and trial structure was used in each case. The staircase procedure estimated the temperature decrease, in the absence of touch, which each participant could detect with a probability of approximately 0.80, called percent-correct point henceforth.

In all experiments, the staircase procedure followed a 3-down/1-up rule. This rule was applied following the first negative response (‘No’). The step sizes were fixed at +0.1°C for the down step and -0.14°C for the up step. The boundaries of the staircase were established at -0.2°C and -2°C. Cooling thresholds of healthy humans lie within this range [11, 26, 27] and the performance of the stimulator was also optimised for this range [11]. The staircase algorithm followed the carry-on rule when the staircase value surpassed the established boundaries [28, 29].

At the start of each staircase trial (figure 2a), the thermal camera started recording to obtain baseline measurements of skin temperature. After 1.5 s, a tone or LED light alerted the participant, and the stimulator shutter opened at the same time, exposing the participant’s skin to the nearby dry ice. When the temperature of the skin in the ROI reached the value assigned by the staircase algorithm, the stimulator shutter closed, the tone or light terminated, and a further beep (duration: 0.2 s; frequency: 100 Hz) indicated that participants should respond. Participants were instructed to answer the same question formulated in the training trials. If the temperature was not reached after a timeout period of 10 s, the trial was considered failed and immediately repeated in another position of the stimulation grid. To refill the stimulator with dry ice and maintain participants’ engagement, there were 2-min breaks every 6-8 min. In all experiments, there were 2 parallel, interleaved staircases: one became progressively colder starting from -0.2°C with respect to baseline skin temperature and the other became progressively less cold starting from -1.2°C. Both staircases were stopped after 12 reversals (figure 2b) [28, 29].

**Figure 2.**
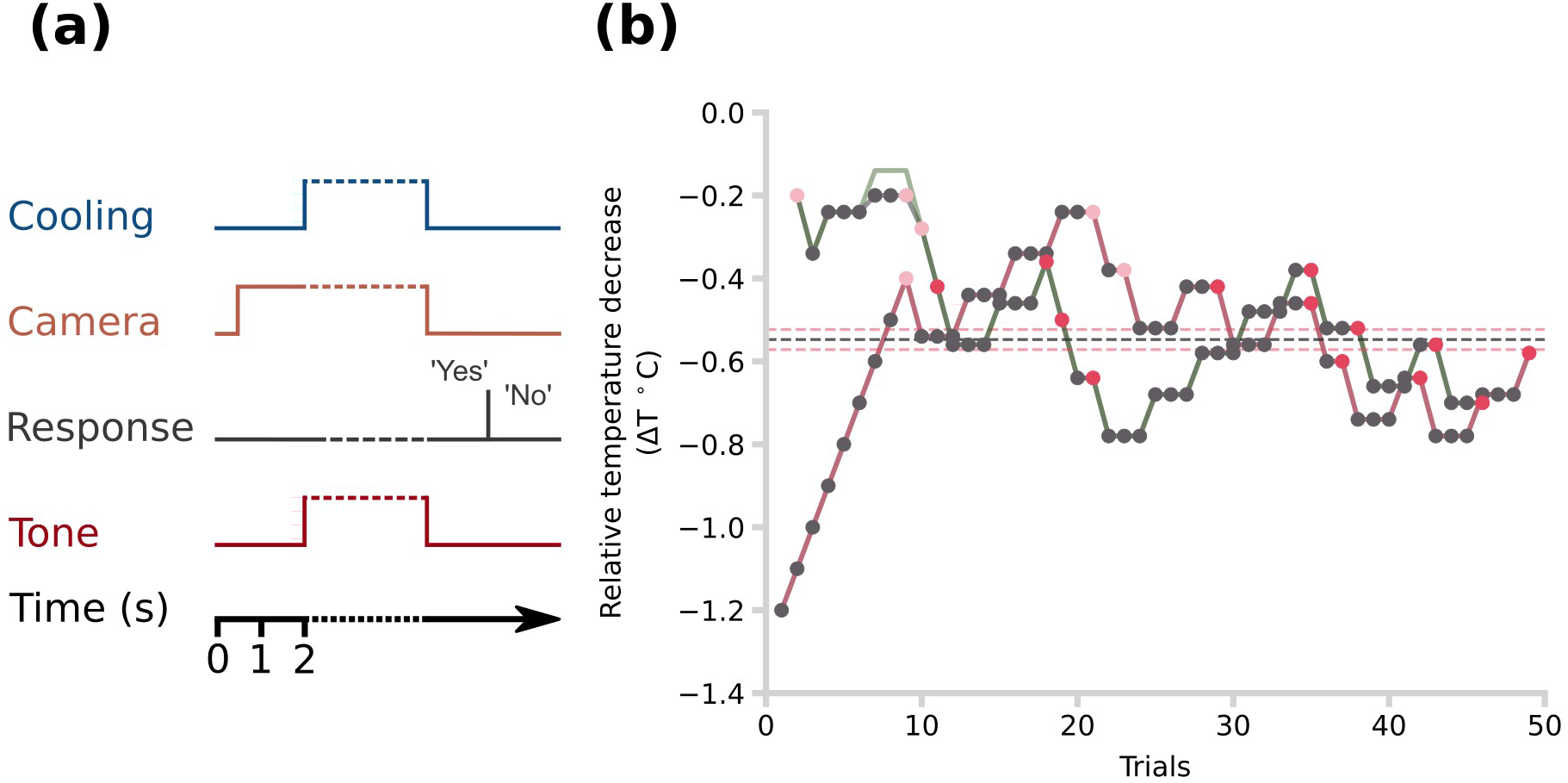
Staircase procedure. **(a)** Schematic of the temporal sequence of events in a trial during the staircase procedure. People responded either ‘Yes’ or ‘No’ to the question: ‘Was there any temperature change during the tone?’ **(b)** An example percent-correct point estimation with a staircase procedure from one participant. The red line follows the value tracked by the staircase algorithm for the descending branch, whereas the green line follows the value tracked for the ascending branch. The black line follows the relative temperature decrease that participants were exposed to at each trial and it is overlaid with the green and red lines for most of the procedure. The black dots indicate the trials in which the participant said ‘Yes’. The light red dots indicate the first three trials in which the participant said ‘No’. These initial trials were excluded. The red dots indicate the subsequent trials in which the participant said ‘No’. The average of these temperatures was taken as the final percent-correct value. The red horizontal dashed lines are the percent-correct points for the descending and ascending staircases. The black line shows the mean of these values.

Experiments 1 and 2 used a signal detection paradigm for two different stimulus types, *Cold* and *Cold & touch*, tested in randomly interleaved order (figure 1b & c). In Experiment 3, there was one signal detection with three conditions: *Cold, Cold & touch* and *Cold & sound*. Each condition consisted of 27 trials in which cooling was present interleaved with 27 trials in which cooling was absent (but other elements of stimulation such as touch and sound were present according to condition). This design allowed us to use signal detection theory [31] (figure 1c) to compare sensitivity and bias of cooling detection with vs. without associated touch or sound.

The structure of trials in the signal detection paradigm was similar in all experiments (figure 1c). First, the thermal imaging acquisition began and the thermal camera took a baseline skin temperature for 0.5 s. Then, for trials involving touch stimulation, the von Frey filaments were moved to touch the skin around the designated cooling stimulation point (figure 1b). For trials involving sound rather than touch, a 500 Hz tone began playing. Next, 2 s later, the shutter of the dry-ice source opened, in cooling (i.e., signal present) trials only (figure 1c). In no-cooling (i.e., signal absent) the shutter was moved to create a comparable noise from the shutter servo-motor, but did not open or expose the source. In experiments 1 and 2, a tone started to alert the participant that cooling might occur. In Experiment 3, the alert was given by an LED, rather than a tone. The thermal camera continually monitored skin temperature in a region of interest under the dry-ice source, and compared this to a baseline measure taken from the first 0.5 s of each trial. Timestamps for individual thermal images showed that this sampling loop operated at 7.24 ± 1.44 Hz. When instantaneous ROI temperature reached the target decrease from baseline estimated as each participant’s 80% detection threshold by the initial staircase (see above), the stimulator shutter closed, the alerting tone terminated (experiments 1 & 2) or the LED light turned off (experiment 3). The duration of each cooling stimulation was recorded and used to replay non-cooling, stimulus absent trials with matched durations. After cooling ended, a brief beep instructed participants to judge whether there had been a temperature change during the tone, exactly as in the initial staircase. Participants either said ‘Yes’ or ‘No’ in Experiments 1 and 2, or pressed a corresponding key in Experiment 3 (figure 1b). The intertrial interval was 8 s. To refill the stimulator with dry ice and maintain the participant’s engagement, there were 2 minute breaks every 6-8 minutes.

As in the initial staircase, failed trials – principally those where the target temperature was not achieved within the 10 s timeout period – were repeated at a random point in the block. Out of 4538 trials, a total of 176 trials (3.7%; mean of 4.9 failed trials per participant) were classified as failed trials. The majority of the failed trials were due to participant movement, which could be corrected immediately after a failed trial thanks to the LED lasers.

### (d) Data Analysis and Statistics

The initial staircase was used to calculate the target temperature change for cooling signal detection in each experiment. The mean temperature change from baseline was estimated from the reversals of each, ignoring the first 3 reversals. A reversal was defined as a trial in which the response of the participant changed relative to the previous trial (figure 2b). The target temperature change values from the interleaved ascending and descending staircases were averaged to produce a final estimate (figure 2b).

For each experiment, the percent correct response, the hit and false alarm rates were calculated for each participant in each condition. The sensitivity to cooling (d’) and the response bias (c) were then calculated using signal detection theory and a standard loglinear method [30-33], which adjusts d’ and C when hit/false alarm rates are 1 or 0. In total, 0% (0/36) of hit rates were 1 and 33% of false alarm rates were 0 (12/36).

We hypothesised that the sensitivity in the ‘Cold & Touch’ condition would be less than the sensitivity in the ‘Cold’ condition, based on Gate Control Theory [1-3]. Therefore, we compared d’ across conditions with one-tailed tests. As we did not have prior predictions about the bias, we compared values of C across conditions with two-tailed paired t-tests. For experiment 3, our predictions focussed on the effects of touch and of sound on cooling detection. Consistent with the previous experiments, the d’ of ‘Cold & touch’ condition was compared to the ‘Cold’ condition with a one-tailed test, whereas the sensitivities and the d’ of the ‘Cold & sound’ condition were compared to the ‘Cold’ condition with two-tailed tests.

## 3. Results

### (a) Touch decreases the sensitivity to focal cooling

The initial staircases for experiment 1 estimated that the smallest temperature decrease from baseline could be detected with 80% accuracy was -0.80°C ± 0.25°C standard deviation. For experiments 2 and 3, the corresponding values were -1.12°C ± 0.54°C and - 1.27°C ± 0.37°C, respectively.

The results of experiment 1 showed that concurrent tactile stimuli (‘Cold & touch’) significantly reduced sensitivity compared to cooling alone (*Cold & touch* d’: 1.25 ± 0.69; *Cold* d’: 1.97 ± 0.66 standard deviation; difference: 0.72 ± 0.52; one-tailed paired-sample t-test; t_11_ = 4.51; p = 0.00004; Cohen’s d = 1.05) (figure 3a). Experiment 2 replicated this result, though with a lower effect size: (*Cold & touch* d’: 1.63 ± 0.85; *Cold* d’: 1.90 ± 0.64; difference: 0.27 ± 0.43; one-tailed paired-sample t-test; t_11_ = 2.09; p = 0.03; d = 0.36) (figure 3c).

**Figure 3.**
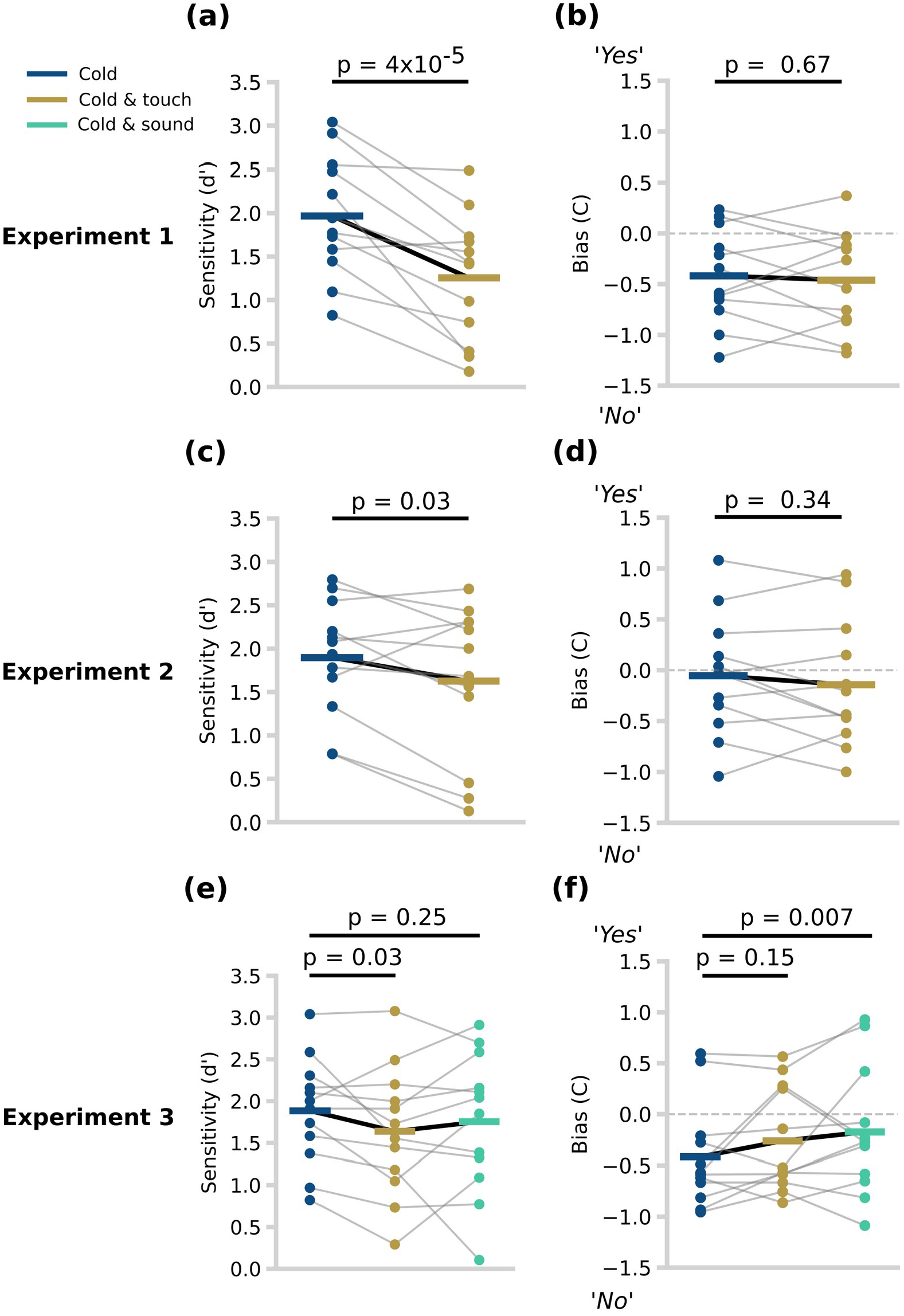
Sensitivity and bias across experiments and conditions. **(a)** The sensitivities (d’) in Experiment 1. Each datapoint (coloured dot) is the sensitivity of each participant during the signal detection paradigm. The light grey lines join the datapoints belonging to the same participant. The horizontal, coloured lines represent the mean of the sensitivities. **(b)** The response biases (C) in Experiment 1. The dashed, horizontal, grey line follows y = 0. A negative value indicates a tendency to say ‘No’, whereas a positive value indicates a tendency to say ‘Yes’. **(c)** The sensitivities (d’) in Experiment 2. **(d)** The response biases (C) in Experiment 2. **(e)** The sensitivities (d’) in Experiment 3. **(f)** The response biases (C) in Experiment 3.

Participants had a tendency to say ‘No’ in both conditions, producing a negative response bias (*Cold & touch*: -0.46 ± 0.47; *Cold* C: -0.42 ± 0.45). There was no significant difference between the two conditions (difference in C: 0.04 ± 0.34; two-tailed paired-sample t-test, t_11_ = 0.43; p = 0.67; d = 0.10) (figure 3b). In experiment 2, participants again had a tendency to say ‘No’ in both conditions (*Cold & touch*: -0.14 ± 0.59; *Cold* C: - 0.05 ± 0.56). There was no significant difference between the two conditions (difference in C: 0.09 ± 0.3; two-tailed paired-sample t-test; t_11_ = 0.99; p = 0.34; d = 0.16) (figure 3d).

### (b) Distraction is unlikely to explain the thermotactile gate

In experiment 3, sensitivity was calculated for each of the three conditions (*Cold* d’: 1.88 ± 0.61; *Cold & touch* d’: 1.64 ± 0.74; *Cold & sound* d’: 1.75 ± 0.80) (figure 3a). Sensitivity was again significantly reduced when non-tactile cooling was accompanied by concurrent tactile stimuli (*Cold & touch* vs *Cold*: difference d’s: 0.25 ± 0.39; one-tailed paired-sample t-test; t_11_ = 2.09; p = 0.03; d = 0.36). There was no significant reduction when non-tactile cooling was accompanied by a sound (*Cold & sound* vs *Cold*: difference d’s: 0.13 ± 0.62; one-tailed paired-sample t-test; t_11_ = 0.70; p = 0.25; d = 0.18)).

Participants had a bias to respond ‘No’ in all three conditions (C values *Cold*: -0.42 ± 0.48; *Cold & touch*: -0.26 ± 0.49; *Cold & sound:* -0.17 ± 0.60) (figure 3b). Planned comparison testing showed no significant effect of concurrent tactile stimuli (*Cold & touch* vs *Cold* difference: -0.16 ± 0.33; two-tailed paired-sample t-test; t_11_ = -1.57; p = 0.15; d = -0.32). However, response bias was significantly changed when non-tactile cooling was accompanied by a sound as compared to the unimodal cooling condition (*Cold & Sound* vs *Cold*: difference: -0.24 ± 0.24; two-tailed paired-sample t-test; t_11_ = -3.33; p = 0.007; d = -0.45). That is, the presence of a sound increased the probability of ‘Yes’ responses, whether cooling stimuli were actually present or not.

Finally, because the *Cold & touch* and *Cold* conditions were present in all three experiments, we additionally performed a planned comparison between these two conditions after pooling across experiments. This confirmed that sensitivity was reduced by touch (*Cold & touch* d’ 1.51 ± 0.79, *Cold d’* 1.92 ± 0.64, difference 0.41 ± 0.50, t(35)=4.85, p<0.001 one-tailed, Cohen’s d=0.57). Pooled analysis of bias showed no significant difference between conditions (*Cold & touch* d’ -0.29 ± 0.54, *Cold d’ -0*.30 ± 0.53, difference 0.01 ± 0.34, t(35)=-0.13, p=0.55 one-tailed, Cohen’s d=-0.01).

## 4. Discussion

We investigated the effect of touch on the detection of focal, non-tactile cooling, using a novel stimulation method that provides non-contact cooling under controlled experimental conditions, and without mechanoreceptor stimulation [11]. Thus, we could measure the sensitivity to focal, non-tactile cooling with and without touch. To our knowledge, this has not been attempted previously. We found that sensitivity to non-tactile cooling was significantly reduced when it was accompanied by touch. Crucially, this effect was specific to mechanoreceptor input, rather than reflecting a general distraction effect of additional stimulation, since detection of cooling was not decreased by a concomitant auditory stimulus balanced for duration and intensity with our tactile stimuli. We suggest our results reflect a previously-overlooked interaction between cooling and tactile signals. We speculate that this interaction may be analogous in its mechanisms and consequences to the well-known interaction between touch and pain described by Gate Control Theory [1-3].

The Gate Control Theory states that non-painful tactile input can suppress pain [1]. Aβ afferent signals are thought to inhibit pain signals carried by Aδ- and C-fibres within the spinal cord, thus reducing the central transmission of the signals that determine perceived pain intensity [2, 3]. Cold sensations are also mediated both by Aδ- and C-fibres [15-16], with Aδ-fibres predominantly responsible for non-noxious cold and C-fibres for noxious cold. A similar gating mechanism may underlie the reduction we observed in sensitivity to non-noxious cooling caused by touch. Specifically, SAI-Aβ fibres activated by static touch may activate inhibitory interneurons, which in turn decrease the transmission of cooling-sensitive Aδ- and C-fibres.

Tactile sensation is mediated by multiple mechanoreceptor types, and their associated afferent fibres, which may function as independent sensory channels or submodalities [4]. Which of these various touch channels might underlie this cold/touch interaction? Green and colleagues previously reported that “dynamic touch” attenuates cold sensation compared to “static touch”. Both of their conditions involved mechanical contact with the thermal stimulator, but the type of contact was quite different. Dynamical touch comprised synchronised changes in both contact force and temperature, for example when a cold object makes new contact with the skin. The static touch condition involved ongoing contact pressure from a stimulator which then changed in temperature [21, 22]. In contrast, our design made thermal and mechanical stimulation completely independent. Further, the transient onset of mechanical contact in Green et al.’s dynamic touch experiments would presumably activate multiple classes of mechanosensitive fibres [34-37]. In contrast, we used two focal mechanical stimuli (i.e. von Frey filaments that 2 s before cooling. Together with previous experiments, our results suggest that the interaction between cooling and tactile inputs might depend on the spatiotemporal profile of mechanical force. Future research should compare the effects of different tactile stimuli on sensitivity to non-tactile cooling.

Spatiotemporal stimulus properties may influence the interaction of thermotactile signals in the nervous system that are not fully understood. Strikingly, sensations of wetness, which are clearly distinguishable from our thermotactile sensations, might emerge from the integration of cooling and tactile signals [6]. For instance, rate of temperature decrease strongly influences wetness perception even in the absence of moisture [38, 39].

A recent study in mice [7] found that the threshold to detect either a cooling or a tactile stimulus decreased when they were presented simultaneously. This might reflect a thermotactile interaction with the opposite sign of the one reported here. There are several differences between these two studies. First, the studies were conducted on different species. Second, the mouse study delivered the thermal stimulus with a contact stimulator, whereas we have used a non-contact stimulator capable of dissociating cooling from mechanical signals. Third, the thermotactile stimuli had different spatiotemporal features. In the mouse study, the tactile stimulus was vibratory and covered the entire dorsal surface of the forepaw throughout the entire experimental session, while the contact thermal stimulus covered the ventral surface of the paw. In our studies, the thermal stimulus had an area of 10.9 mm^2^ [11] and was delivered to the dorsal surface of the hand. The tactile stimuli bracketed the thermal stimulus, and had a diameter of 0.503 mm^2^. Therefore, the difference in the direction of the effect could be due to differences in the cooling and tactile stimuli. Future research should study the mechanism underlying differences in perceptual output across stimuli space as this might reveal overlooked receptors, fibre types and pathways. For example, the suppressive effect of touch on cold sensitivity that we have found should be investigated with parametric variations of the both thermal and tactile stimulus area.

The brain has limited resources for processing sensory information. Therefore, it could be that touch is simply a distraction for detecting cooling and the effect we observe is due to attentional shift rather than to a gating mechanism. In our study, we minimised attentional effects in four ways. First, the tactile stimulus was never relevant to the task. Second, in all trials there was either a tone or a light that alerted the participant when temperature changes might occur. Temporal expectancy was therefore balanced across conditions and independent of the presence of touch. Third, our tactile stimulus was designed to avoid shifts in *spatial* attention, since the two monofilament stimuli were centred on the cooling location. Finally, the filaments always touched the skin 2 s before the onset of cooling and then remained static until the end of cooling. New events attract attention transiently (“exogenous attention”) for around 200 ms [40], but sustained stimuli may not attract attention (e.g., we tend to ignore tactile input from our clothes).

Further, experiment 3 included a condition with an auditory stimulus to control for attentional, arousal and distraction effects of multisensory stimulation. We found no evidence that the concurrent sound modulated sensitivity to cooling, though we found that the sound did induce a shift towards more liberal response bias. In contrast, concomitant touch did not significantly influence response bias in any experiments. Some participants in experiment 3 spontaneously volunteered that they had found difficult to stay alert and engaged on trials without a tone. We therefore speculate that the tone may have had attentional effects. Since experiments 1 and 2 included a tone on all trials, the effects of touch on cooling detection would be independent of any such attentional effects. Further, we found that touch inhibited sensitivity to cooling across all three experiments, despite differences in other aspects of the trial structure, such as the alerting signals used. Therefore, it seems unlikely the inhibitory effect of touch on sensitivity to cooling we found is due to attentional mechanisms.

We note some limitations of our methods and results. First, we cannot know exactly what classes of afferents are activated by our dry-ice cooling, nor by our monofilament tactile stimulation. The hypothesised inhibitory interaction between tactile and thermal signals has not been confirmed directly by neurophysiological data. Our hypothesis that Aβ fibres interneuronally inhibit transmission of signals by Aδ fibres therefore remains speculative. Future microneurographic studies could attempt to record from individual afferents of these classes during stimulation using our experimental conditions, and then relate behavioural effects to firing patterns. However, microneurography is limited to opportunistic sampling from peripheral afferents, so cannot reliably identify changes in afferent signals due to spinal interactions. Animal studies could successfully study spinal interactions between specific signals [41], but present limitations for studying conscious experience.

Second, we cannot completely exclude some incidental mechanical effect of dry-ice cooling, due to convection currents. We measured the force on the skin generated by downward airflow through our cooling apparatus at 0.53 mN [11]. This is below published threshold values for activating slowly adapting SAI and SAII units (1.3 mN and 7.5 mN, respectively) [42, 43], suggesting the forces generated by convection are negligible. Further, any mechanical effect from dry-ice thermal sensation should be similar in all our experimental conditions, so cannot readily explain differences between touch-present and touch-absent conditions. Third, while the inhibitory effect of touch on sensitivity to cooling was present across all three experiments, it varied somewhat in size. The reasons for this variation are not clear. The three experiments were performed in two different laboratory rooms, and at two different seasons, so contextual factors might have contributed to variability in effect size. Future, larger studies might provide a more stable estimate of mean effect size, and a clearer picture of why the effect size may vary across individuals.

In conclusion, we report an apparently novel interaction in thermotactile somatosenation. Specifically, touch reduces detection sensitivity for focal, non-tactile cooling. Classic views of cortical somatosensation suggest that signals for each submodality ascend independently to primary cortex. Only then, in secondary and associative cortical regions, is somatosensory information integrated across different submodalities to produce an overall percept [44, 45]. These cortical interactions are often linked to causal inference computations [46], and to a general prior of objects having parallel multisensory attributes [47]. An alternative view suggests that perception is shaped by multiple interactions between afferent signals at each step along the ascending somatosensory pathway. In particular, elaborate patterns of interaction in the spinal cord can be identified by anatomical studies [4, 5], potentially explaining the robust finding of tactile gating of nociceptive afferent signalling, leading to reduced pain levels [1-3]. Our findings add a novel interaction between touch and temperature to this interaction-based view, and contribute to our understanding of inter-channel interactions in somatosensation. Our study could also lead to potential applications in areas such as clothing design, and wearable technology. Further perceptual and neurophysiological studies are required to confirm the precise neural mechanism of the interaction we have identified.

## Data accessibility

The data shown in this manuscript and the code for collecting, analysing and visualising it can be found in the following link: https://github.com/iezqrom/publication-touch-inhibits-cold. More information around the non-tactile cooling stimulator including additional data and code can be found in a previous study [11].

## Declaration of AI use

We have not used AI-assisted technologies in creating this article.

## Authors’ contributions

I.E.R.: data curation, formal analysis, funding acquisition, investigation, methodology, software, validation, visualization, writing—original draft, writing—review and editing; M.C.: investigation, methodology, writing—review and editing; P.H.: conceptualization, funding acquisition, resources, methodology, project administration, validation, supervision, writing—review and editing.

## Conflict of interest declaration

We declare we have no competing interests.

## Funding

The Biotechnology and Biological Sciences Research Council (UK) [grant number BB/M009513/1] supported I.E.R. The European Union Horizon 2020 Research and Innovation 385 Programme, TOUCHLESS (project No. 101017746) supported M.C. & P.H..

## Acknowledgements

The authors would like to thank Martin Donovan for his technical support in the development of the methodology. Help from Dr. Shinya Takamuku was also greatly appreciated. Informal discussions with Angel Ezquerra and Guanhaven Romano-Mendoza were helpful in the inception and development of the methodology.

## Notes

https://github.com/iezqrom/publication-touch-inhibits-cold

### Competing Interest Statement

The authors have declared no competing interest.

